# Regulation of protein complex partners as a compensatory mechanism in aneuploid tumors

**DOI:** 10.1101/2021.12.05.471308

**Authors:** Gökçe Senger, Stefano Santaguida, Martin H. Schaefer

## Abstract

Aneuploidy, a state of chromosome imbalance, is a hallmark of human tumors, but its role in cancer still remains to be fully elucidated. To understand the consequences of whole chromosome-level aneuploidies on the proteome, we integrated aneuploidy, transcriptomic and proteomic data from hundreds of TCGA/CPTAC tumor samples. We found a surprisingly large number of expression changes happened on other, non-aneuploid chromosomes. Moreover, we identified an association between those changes and co-complex members of proteins from aneuploid chromosomes. This co-abundance association is tightly regulated for aggregation-prone aneuploid proteins and those involved in a smaller number of complexes. On the other hand, we observe that complexes of the cellular core machinery are under functional selection to maintain their stoichiometric balance in aneuploid tumors. Ultimately, we provide evidence that those compensatory and functional maintenance mechanisms are established through post-transcriptional control and that the degree of success of a tumor to deal with aneuploidy-induced stoichiometric imbalance impacts the activation of cellular protein degradation programs and patient survival.

## Introduction

Aneuploidy, whole chromosomal or chromosome arm-level alterations, affects expression of a large number of genes simultaneously - most directly by providing additional or diminished copies of genes in- or decreasing their transcription. Typically, aneuploidy is detrimental for normal cells, among other reasons as it causes stoichiometric imbalances in protein complexes involving proteins encoded on the aneuploid chromosome^1,2^. However, around 90% of solid tumours have aneuploid karyotypes^3,4^, raising the question of how cancer cells can tolerate the massive amount of transcriptomic and proteomic changes.

Previous studies demonstrated correlated expression between gene copy numbers and transcriptome level for genes on aneuploid chromosomes while there is a buffering effect at the proteome level adjusting protein levels, especially for protein complex subunits^5,6^. This post-transcriptional compensatory mechanism preventing excess translation of complex members has also been characterized in the context of copy number alterations (CNAs) in different contexts such as yeast^7^ and cancer^8^. Structural properties of proteins have an effect on the degree of post-transcriptional buffering on changes induced by CNAs, for example proteins with larger interface size showed larger degree of buffering^9^. Recently, post-transcription regulation has been identified as a dosage compensation mechanism in response to aneuploidy in cancer cell lines^10^.

Transcriptome analysis revealed that aneuploidy largely affects expression of genes on other chromosomes too^11,12^. For CNAs a correlated increase in the abundance of co-complex members encoded by genes outside the copy number amplified region has been shown^8^ raising the question of whether this could contribute to the expression changes in aneuploid cells even on diploid chromosomes. Together these results suggest the importance of compensation mechanisms to buffer differentially expressed transcripts in response to the amplification or loss of genomic regions and in particular to mitigate stoichiometric imbalances in protein complexes. However, we still lack a global understanding on the effects of aneuploidy on the expression of genes on other, non-aneuploid chromosomes in a cancer context.

Here, we study transcriptomic and proteomic changes induced by aneuploidy in cancer patients (Figure 1A) and extend the scope of previous studies by focusing on expression and abundance changes on other, non-aneuploid chromosomes. We show that protein complex subunits of other chromosomes tend to maintain their abundance levels unless they form a complex with differentially abundant proteins encoded by genes located on the aneuploid chromosome. We further demonstrate that this co-abundance regulation is dependent on aggregation propensity and promiscuity of the aneuploid complex partners and controlled by post-transcriptional mechanisms. Our findings highlight a complementary mechanism acting to deal with the excess amount of expression changes induced by aneuploidy: Coordinated abundance changes of complex partners might prevent aggregation of unpaired complex members.

**Figure 1 with 1 supplement.**
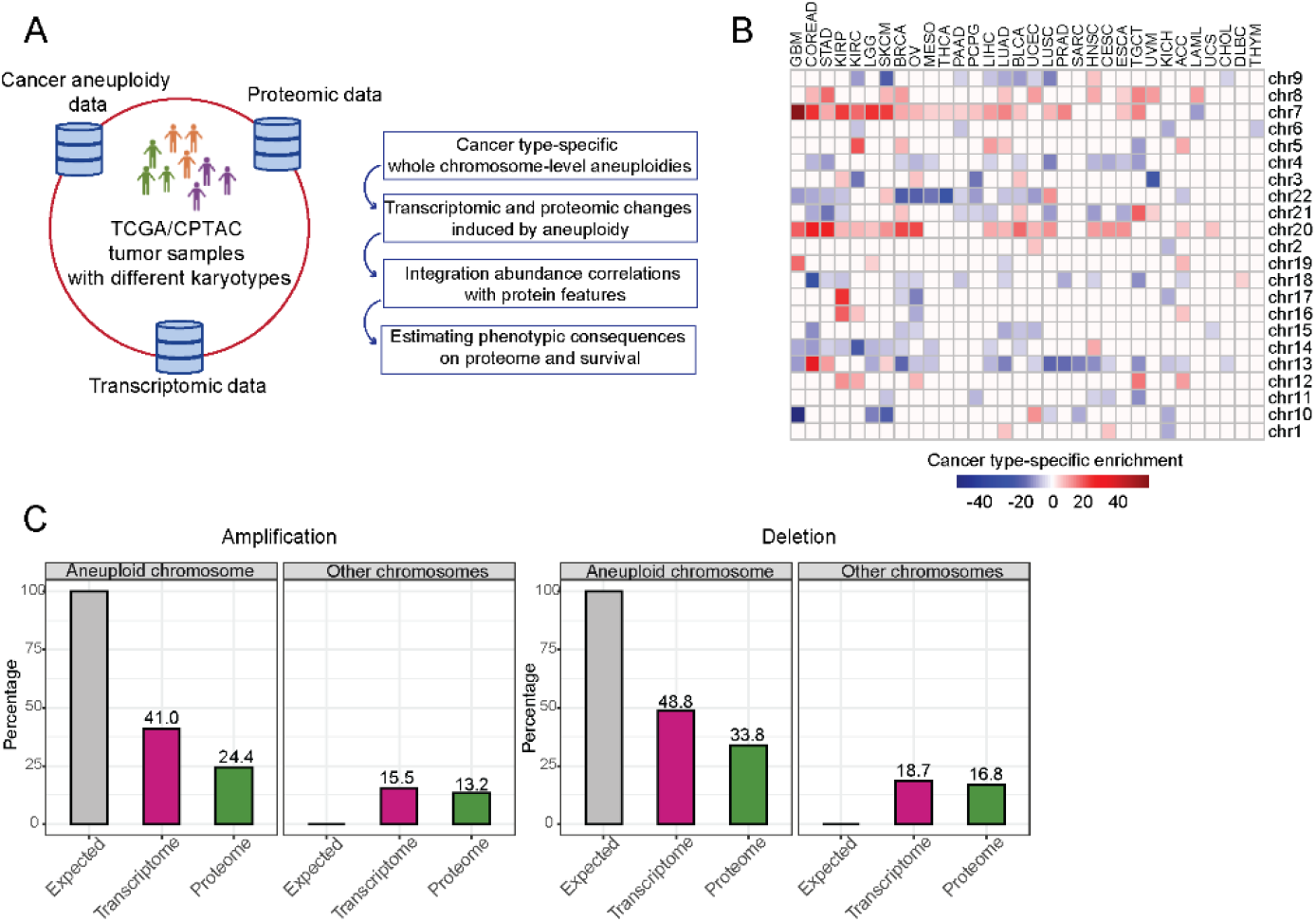
Transcriptomic and proteomic changes in aneuploid tumors. **A)** Data used in this study and schematic representation of the performed analyses. **B)** Cancer type-specific whole chromosome-level alterations across 32 cancer types. The color encodes the degree of their enrichment (standard residuals of the chi-square test multiplied by the alteration score [-1 in the case of deletion and 1 in the case of amplifications]). **C)** Average percentage of differentially expressed genes or abundant proteins on aneuploid and other, non-aneuploid chromosomes (among the detected genes on the respective chromosomes).

## Results

### Widespread transcriptome and proteome deregulation in aneuploid tumors

To study the effect of whole chromosomal alterations on cancer transcriptomes and proteomes, we first identified cancer-type specific whole chromosome-level amplifications and deletions that occurred at higher frequencies than would be expected by chance in 10522 samples analyzed in The Cancer Genome Atlas (TCGA) by using a previously established cancer aneuploidy estimate^4^. In total, we detected 203 whole chromosome-level alterations including 86 amplifications and 117 deletions for 32 cancer types (Figure 1B, Supplementary file 1).

Then, for each detected aneuploidy case, we split the set of samples into those containing the respective chromosome number aberration and those diploid for the respective chromosome and then tested for differential expression of all genes between the sets. We found that on average 41% and 48% of the genes located on amplified and deleted chromosomes, respectively, changed expression (Figure 1C). Besides those intuitively expected gene expression changes on the aneuploid chromosomes, we observed a surprisingly large number of expression changes happening on other, typically diploid chromosomes (15% and 18% of genes on average for amplification and deletion cases, respectively; Figure 1C).

We observed that often chromosomes tend to be co-amplified. We therefore tested for statistical dependence between amplification events of chromosomes. For the 86 cancer-type specific amplifications, we identified 305 co-amplifications that occurred more frequently than expected by chance (adjusted p-value < 0.01, chi-square test; Supplementary file 1). We wondered if this could explain the relatively high number of differentially expressed genes on other chromosomes. To test this, we identified the strongest case with the smallest p-value (chromosome 7 in thyroid cancer [THCA]) and determined the differentially expressed genes between THCA samples with chromosome 7 amplified vs chromosome 7 diploid. We then quantified the number of differentially expressed genes from each chromosome normalized by the total number of genes on that chromosome (Figure 1 - figure supplement 1B). We found that the number of differentially expressed genes from the co-amplified chromosomes (5, 16 and 20) were on the lower end of the distribution suggesting that they do not substantially contribute to the overall transcriptional dysregulation in chromosome 7-amplified samples.

To further understand the effect of those expression changes in response to aneuploidy on the proteome, we collected corresponding proteome abundance data, which is available from the Clinical Proteomic Tumor Analysis Consortium (CPTAC) for 298 TCGA tumour samples comprising breast (BRCA)^13,14^, ovarian (OV)^15,16^, and colorectal adenocarcinoma (COREAD)^17,18^ cancer types. Then, for 13 and 20 cancer-type specific whole chromosome-level amplifications and deletions, respectively (found in above mentioned 3 cancer types), we detected protein abundance changes between aneuploid samples (with the amplified/deleted chromosome) and diploid samples. We observed that a smaller number of proteins showed abundance changes compared to our observations at the transcriptome level (after normalizing for the largely different gene coverage between the transcriptomics and proteomics datasets) both for amplification and deletion cases (Figure 1C).

Together these observations suggest that attenuation mechanisms are in place, and we observe regulation both on transcription and translation level which prevent that all genes located on the aneuploid chromosomes are deregulated. This effect is stronger on protein than on RNA level indicating that on top of regulation of transcription, translation control or protein degradation might play a role. At the same time, we observed a surprisingly large number of dysregulation events on chromosomes other than the aneuploid one raising the question of the purpose of the up- and downregulation of hundreds of genes in response to specific aneuploidies.

### Complex members tend to be co-deregulated

To test if the vast changes on the cellular proteome in aneuploid cells outside the aneuploid chromosome could be explained by compensation mechanisms for changes in complex stoichiometry induced by aneuploidy, we performed an association test between co-complex members of differentially abundant proteins encoded on aneuploid chromosomes and differentially abundant proteins encoded on other chromosomes by using human protein complex information from the mammalian protein complex database CORUM^19^. We observed a general tendency for the differentially abundant proteins of other chromosomes to be complex partners of differentially abundant proteins from the aneuploid chromosome for both whole chromosome-level amplifications and deletions (p-value < 0.05, chi-square test; Figure 2A). We found a moderate percentage (in average is 4.47%; Supplementary file 2) of differentially abundant proteins on other chromosomes being partners of those on aneuploid proteins.

**Figure 2 with 1 supplement.**
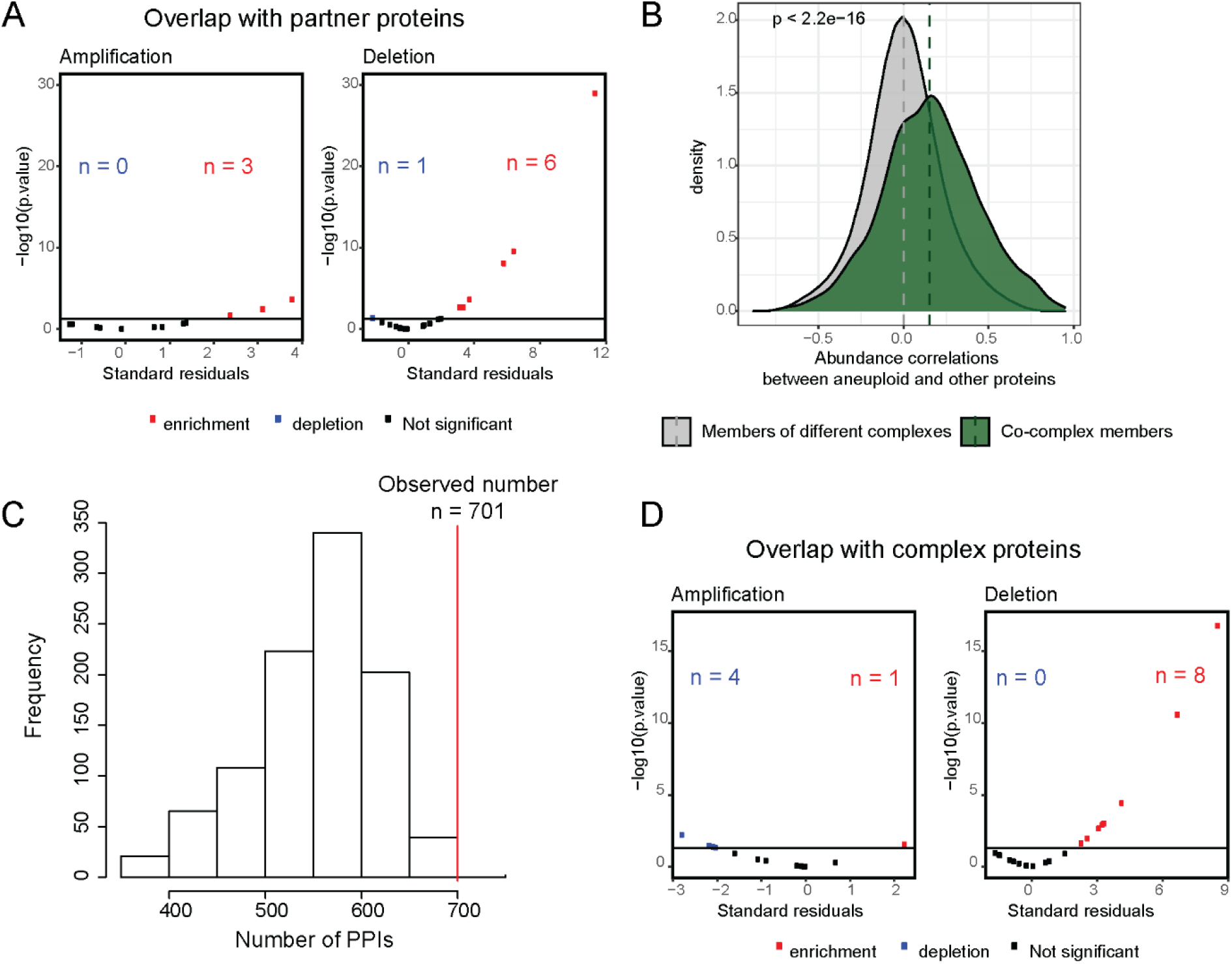
Enrichment of partners of aneuploid proteins in differentially abundant proteins on other chromosomes. **A)** Standard residuals and p-values for the overlap between co-complex members of differentially abundant proteins on aneuploid chromosomes and differentially abundant proteins on other chromosomes for 13 amplifications and 20 deletions. **B)** Protein abundance correlations between differentially abundant proteins on aneuploid chromosomes and their co-complex and non-complex subunits. Correlations were calculated across cancer samples, separately for each cancer type, and then pooled. Wilcoxon test was used to determine whether two distributions are significantly different. **C)** The number of PPIs (n = 701) between differentially abundant proteins of aneuploid chromosomes and those on other chromosomes against the background distribution for COREAD chromosome 7 amplification. **D)** Standard residuals and p-values for the overlap between CORUM complex subunits and differentially abundant proteins on other chromosomes for 13 amplifications and 20 deletions.

However, the coverage of proteins with CORUM complex information is rather limited (22% of proteins form part of at least one complex in CORUM). When only proteins participating in at least one complex were considered, the average fraction of partner proteins among differentially abundant proteins increased to 12.61% (Supplementary file 2). Moreover, comparing protein abundance correlations between differentially abundant proteins on aneuploid chromosomes and their co-complex members with non-complex members showed significantly stronger correlations between proteins of same complexes (p-value < 2.2e-16, Wilcoxon test; Figure 2B). Those observations are in line with previous findings claiming that complex organization shapes protein abundance changes in response to CNAs^9^.

To investigate whether our observations can be generalized to binary protein-protein interactions (PPIs), we used PPI data from the human protein-protein interaction database HIPPIE (v2.2)^20^ to test if the number of interactions between differentially abundant proteins encoded on the aneuploid and those on other chromosomes is higher than expected by chance. Indeed, we found an enrichment of interactions between those protein sets in 9 out of 13 cancer type-specific amplifications and 8 of the 20 deletions (p-value < 0.05, randomization test; Figure 2C, Supplementary file 3). Given the higher coverage of PPI data, we asked again which percentage of differentially abundant proteins on other chromosomes could be potentially explained by their interactions with complex members on aneuploid chromosomes. We found that on average 27.5% of the differentially abundant proteins on other chromosomes interact with those on the aneuploid chromosomes (Supplementary file 3). For chromosome 7 in COREAD and chromosome 12 in OV, more than 40% of the differentially abundant proteins interacted with proteins on the amplified chromosomes.

We hypothesized that these abundance changes should only affect co-complex members of differentially abundant proteins on aneuploid chromosomes, but that non-partner complex members should maintain their abundance level to prevent stoichiometric imbalances. To test this, we performed an association test for the overlap between differentially abundant proteins on other chromosomes and all known human complex members curated from CORUM. As a result of this, we found a significant depletion of complex subunits in differentially abundant proteins on other chromosomes for amplification cases (p-value < 0.05, chi-square test; Figure 2D). This suggests that complex members overall are stably expressed to prevent disruption of complex stoichiometry upon chromosomal amplification. This effect was not observed in the case of chromosomal deletions in which differentially abundant proteins of other chromosomes are significantly enriched in complex proteins (p-value < 0.05, chi-square test; Figure 2D).

In contrast to our observations at the proteome level, we observed both strong enrichments and depletions of co-complex members of proteins encoded by differentially expressed genes on aneuploid chromosomes in the differentially expressed genes on other chromosomes (adjusted p-value < 0.01, chi-square test; Figure 2 - figure supplement 1A). In addition, we observed a significant enrichment of protein complex subunits among the differentially expressed genes on other chromosomes for 19 out of 28, and 20 out of 31 significant associations for amplification and deletion cases, respectively, at the transcriptome level (adjusted p-value < 0.01, chi-square test; Figure 2 - figure supplement 1B). These results together with consistent partner associations that we observed at the proteome level suggest post-transcriptional compensatory mechanisms to control abundance changes induced by aneuploidy.

### Differential protein abundance of complex partners as a compensatory mechanism to prevent complex imbalance and aggregation

The main expected detrimental effect of chromosomal amplifications is an excess of protein abundance of complex members leading to an aggregation of the orphan proteins (rather than a loss of function of the complex as would be expected for insufficient expression for complex assembly as a consequence of chromosome deletion)^21^. We therefore tested if aggregation-prone proteins on the amplified chromosome show a higher tendency for strong correlations with their complex partners on other chromosomes. We grouped aneuploid proteins as aggregating and non-aggregating based on the data from Maata et al^22^ and compared their protein level abundance correlations with partners. Indeed, we observed stronger correlations for aggregation-prone proteins as compared to their non-aggregating counterparts (p-value < 2.2e-16, Wilcoxon test; Figure 3A). This suggests that upregulating the protein expression of genes on chromosomes not affected by aneuploidy themselves serves as a compensatory mechanism to prevent proteotoxicity triggered by the aggregation of non-paired complex members located on the aneuploid chromosome.

**Figure 3.**
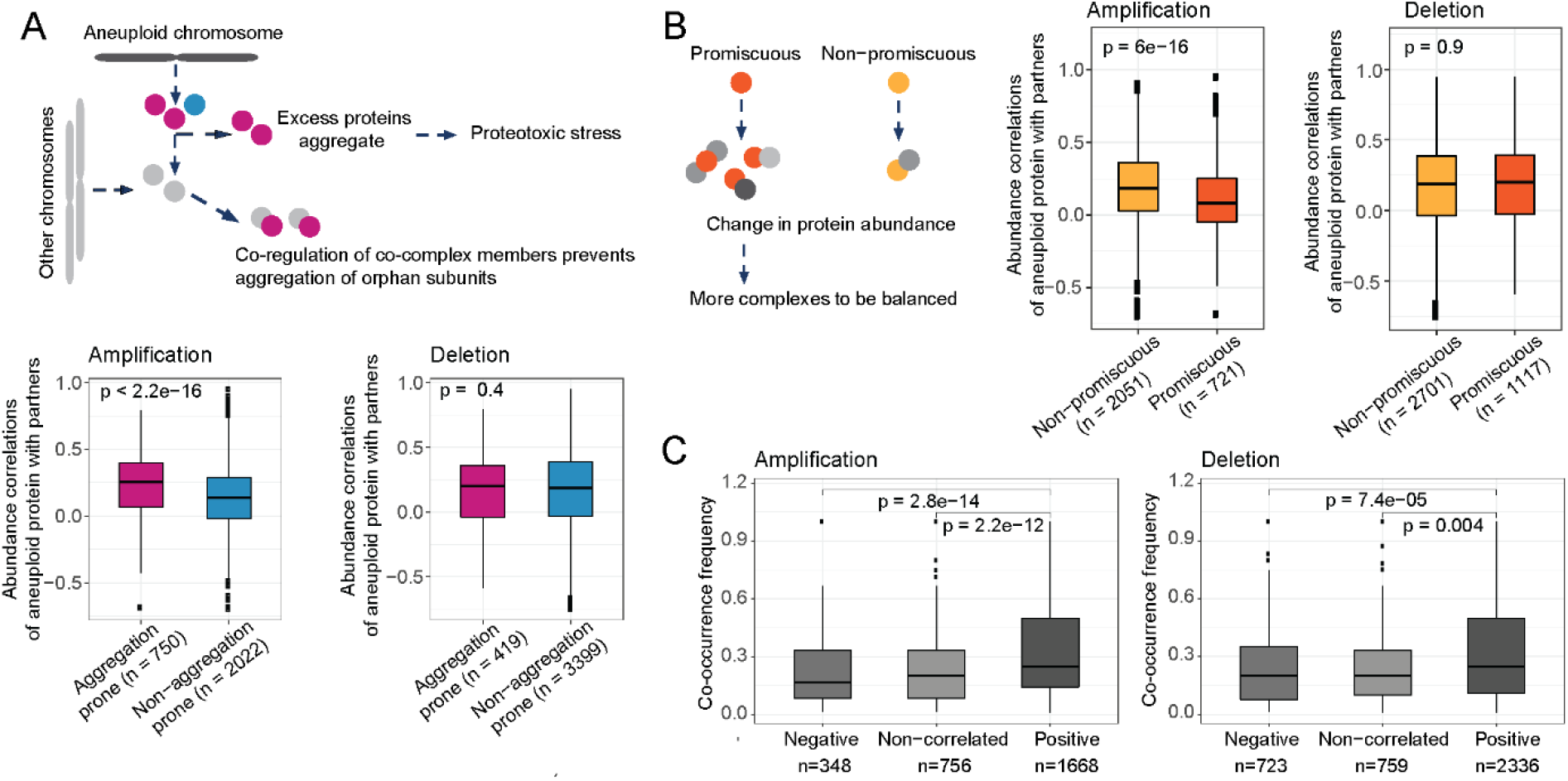
Compensatory mechanisms preventing aggregation and imbalances in complexes partly explain abundance changes on other chromosomes. Protein abundance correlations between differentially abundant proteins on aneuploid chromosomes and their co-complex members on other chromosomes when aneuploid proteins are grouped as **A)** aggregation prone and non-aggregation prone and **B)** promiscuous and non-promiscuous. **C)** Co-occurrence frequency of differentially abundant proteins on aneuploid chromosomes and their co-complex members on other chromosomes in different correlation groups, positively- and negatively-correlated and non-correlated pairs. Wilcoxon test was used to test differences between groups.

We hypothesized that in the case of chromosomal deletions, the aggregation propensity of downregulated proteins on the aneuploid chromosome should not affect the degree of correlation with complex partners. Indeed, we observed aggregation-prone proteins to be not related to stronger correlations with their complex partners on other chromosomes when they are encoded on deleted chromosomes (Figure 3A). This is likely the case as downregulating those proteins would not leave them as orphan subunits and hence increase their risk of aggregation.

Assuming that the regulation of proteins on other, non-aneuploid chromosomes serves to prevent stoichiometric imbalance of protein complexes, we speculated that for proteins that are in many complexes there are more ways of being abundance-compensated by a complex partner compared to those proteins participating in few complexes and therefore each single partner should be under less stringent control for coexpression with the aneuploid protein. We therefore classified each aneuploid protein into promiscuous (participating in more than 5 complexes) and non-promiscuous (involved only in 5 or less than 5 complexes). As expected, we observed weaker correlations for promiscuous proteins of amplified chromosomes (p-value = 6e-16, Wilcoxon test; Figure 3B) further supporting the model in which differential abundance of proteins on other chromosomes is a compensatory mechanism. We did not observe the same association in the case of chromosomal deletions (Figure 3B).

Finally, we hypothesized that proteins co-occurring in many complexes should show stronger correlation as proteins found only in a few cases together in the same complex. Indeed, we found significant differences in the number of times positively correlated protein pairs were found together in the same complex vs uncorrelated or negatively correlated protein pairs (p-value = 2.2e-12 and p-value = 2.8e-14 for chromosomal amplifications, p-value = 0.004 and p-value = 7.4e-05 for deletions; Figure 3C). This, again, illustrates how complex organization shapes the co-abundance patterns between differentially abundant proteins from the aneuploid chromosome and those located on other chromosomes.

### Functional selection acting on protein stoichiometric imbalance

In the previous sections, we proposed that co-abundance changes of protein complex partners is a compensation mechanism to prevent stoichiometric imbalance in protein complexes to avoid proteotoxicity of orphan subunits. We next wondered if besides biophysical (such as aggregation propensity) any functional properties would protect complexes and complex subunits from abundance imbalance in aneuploid cancer cells. To this end, we first retrieved the most strongly correlated co-complex members of differentially abundant aneuploid proteins and identified the complexes they are involved in. Then, we obtained functional annotations of the complexes from the CORUM database. To identify functions under stronger protection from protein abundance imbalance in complexes, we computed the enrichment of these functions compared to a random set of complexes under relaxed stoichiometric protection. The analysis revealed that top correlated proteins form complexes that are frequently involved in translation, RNA splicing, RNA processing and protein complex assembly (Figure 4A; Supplementary file 5). Interestingly, the functional enrichment is consistent for amplifications and deletions suggesting that not just compensatory mechanisms to prevent proteotoxicity contribute to the dysregulation of proteins on other chromosomes but also functional selection is in place, acting on important cancer-essential functions up- or downregulating entire protein complexes while keeping their stoichiometry in check.

**Figure 4.**
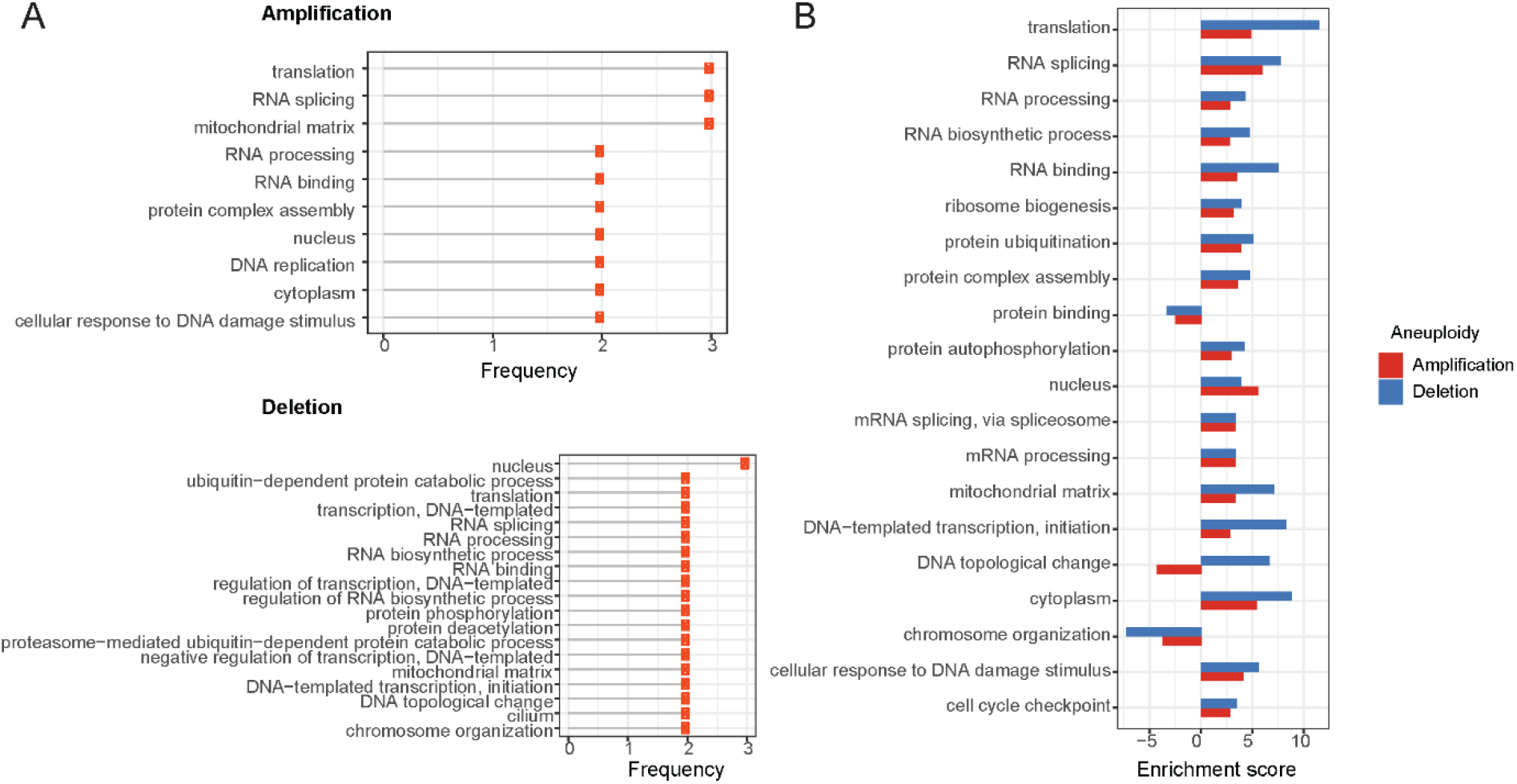
Enrichment of functional terms in complexes of top correlated proteins. **A)** Most frequently enriched terms in the amplification and deletion cases. Frequency shows the number of aneuploidy cases in which the corresponding term is significantly enriched. **B)** Enrichment scores of enriched terms both in amplification and deletion cases. For the functional term that is enriched in more than one amplification/deletion cases, the enrichment score of the ones with the lowest p-value is displayed.

To quantitatively compare the degree of enrichment between the functional terms associated with balance-protected complexes, we devised an enrichment score (see Methods section) and compared it for the top enriched or depleted functions between amplifications and deletions. We observed that top correlations in the deletion cases are related to stronger enrichment scores when compared to their counterparts in the amplification cases (Figure 4B; Supplementary file 5).

### Post-transcriptional regulation of partner co-abundance

Our results suggest a central role for post-transcriptional regulation in maintaining complex protein abundance balance in aneuploid cells. We therefore aimed to further disentangle the different regulatory layers underlying expression changes (Figure 5A) on other chromosomes induced by aneuploidy. First, we tested if those expression changes could be explained by epigenetic silencing via differential DNA methylation of the promoters of genes changing expression in aneuploid samples. Therefore, we compared average DNA methylation levels of differentially expressed genes in aneuploid samples to diploid samples, separately for up- and down-regulated genes. We found that down-regulated genes are significantly related to higher methylation levels in aneuploid samples in only 6 amplification cases out of 86 (∼7%) and in 5 deletion cases out of 117 (∼4%; Figure 5 – figure supplement 1A). We observed significant associations between lower methylation level and up-regulated genes in aneuploid samples for few cases (10 out of 86 amplification cases and 16 out of 117 deletion cases) (Figure 5 – figure supplement 1B). This suggests that epigenetic regulation does not have a substantial contribution to the described genome-wide changes in gene expression.

**Figure 5 with 1 supplement.**
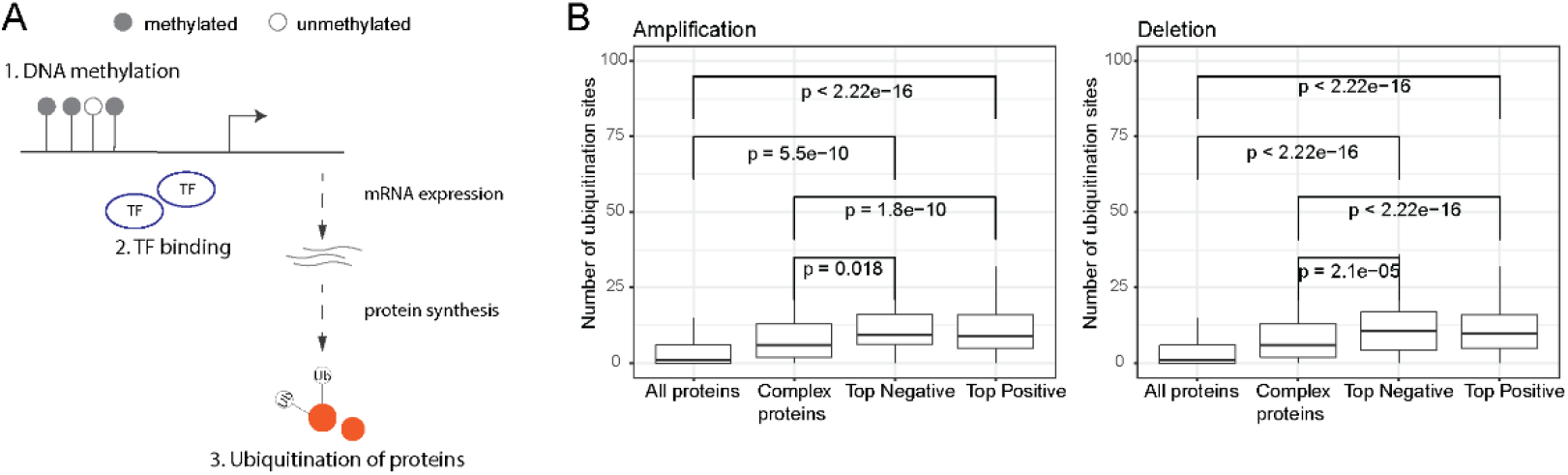
Post-transcriptional regulation of co-complex members of aneuploid proteins. **A)** Overall representation of different levels of gene regulation. **B)** Number of ubiquitination sites of all, human complex, and top positively and negatively correlated proteins. Wilcoxon test was used to test differences between groups.

We then asked if differential expression of transcription factors (TFs) on the aneuploid chromosome could explain the large transcriptional changes on other chromosomes. We therefore tested for a large list of ENCODE gene-TF associations^23,24^, if there is an enrichment of targets of differentially expressed TFs on the aneuploid chromosome among differentially expressed genes on the other chromosomes. Performing a randomization test did not reveal an excess of targets for any of the tested, cancer type-specific chromosomes (Figure 5 – figure supplement 1C). Together with the methylation analysis, these results suggest that transcriptional regulation cannot fully explain the expression changes on other chromosomes.

As we observed a stronger association between complex partner co-regulation on proteome as compared to transcriptome-level, we hypothesized that post-translational modifications could regulate co-abundance changes of partners of aneuploid proteins on other chromosomes. To test this, we retrieved ubiquitination data from PhosphoSitePlus^25^ and tested if top-correlated co-complex members of aneuploid proteins tend to have higher number of ubiquitination sites (as a proxy to identify proteins that can be more easily targeted for degradation). Indeed we found that top correlated partners have significantly higher numbers of ubiquitination sites (p-value < 0.05, Wilcoxon test; Figure 5B). This suggests that a primary mechanism for keeping protein complex stoichiometry in check seems to be indeed post-transcriptional regulation (such as ubiquitin-mediated degradation).

### Phenotypic consequences of stoichiometric compensation success

The previous results suggest co-regulation of co-complex members as a compensation mechanism to balance protein abundance changes caused by whole chromosomal alterations and thus to keep protein complex stoichiometry in check. We reasoned that different tumors might be able to compensate for the dysregulation of proteins on the aneuploid chromosome with a different degree of success and hypothesized that tumors that can better compensate for protein abundance changes will be associated with better survival rates while those that fail to compensate should upregulate components of the protein degradation machinery to clear the cell from the orphan complex subunits. To test this, we first calculated a stoichiometry deviation score for each sample by using correlations between co-complex members in aneuploid tumors as a measure of failure of keeping the complex stoichiometry balance (Figure 6A). Then, we performed a survival analysis by grouping samples based on their stoichiometry deviation scores (Figure 6A). While not significant in every single case we observed a tendency that samples with low stoichiometry deviation scores are related to lower survival probabilities in all three tissue types (Figure 6B) showing that compensation for protein abundance indeed provides a survival advantage to tumors.

**Figure 6 with 1 supplement.**
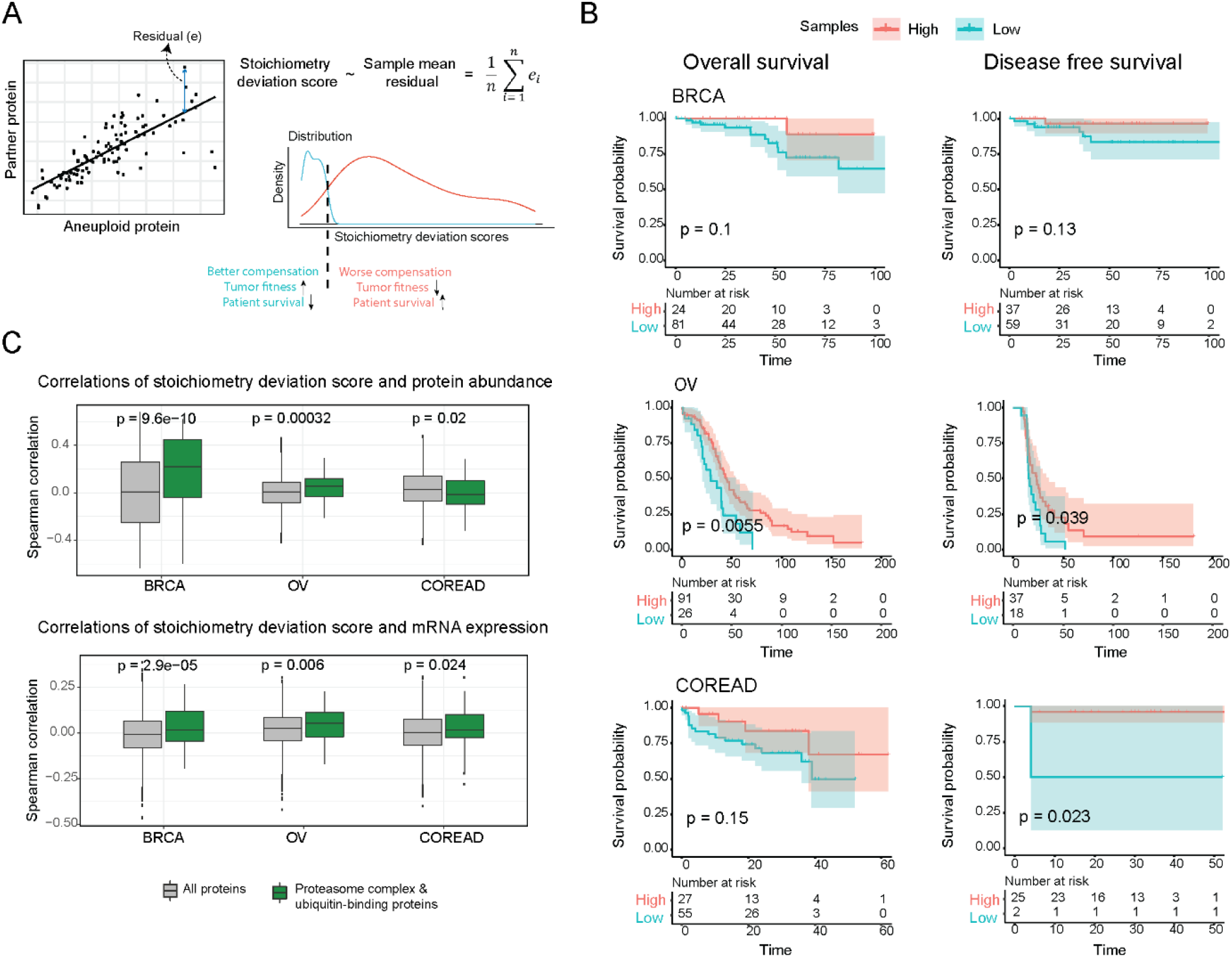
Consequences of stoichiometric compensation. **A)** Graphical representation for the calculation of stoichiometric deviation score for each sample. **B)** Survival analysis results within each tissue. Survival analysis was done once with overall survival and once with disease free survival. **C)** Correlations between the stoichiometric deviation scores and protein abundance/mRNA expression of all proteins, and proteasome complex - ubiquitin binding proteins. Wilcoxon test was used to test differences between groups.

We further investigated if the proteins that play a role in protein degradation have higher abundances in the tumors that cannot compensate for abundance changes and thus have to deal with the excess amount of orphan subunits. We indeed found that ubiquitin-binding proteins and components of the proteasome show significantly higher correlations between their abundances and the stoichiometry deviation scores in two out of three tissues (Figure 6C; Figure 6 – figure supplement 1B). This likely is a consequence of proteotoxic stress resulting from the inability of some tumors to keep protein complexes balanced.

## Discussion

Here, we conduct an extensive characterization of the transcriptome and proteome in aneuploid human cancers. Consistent with previous results^5,6^, we find that proteins on aneuploid chromosomes tend to be buffered in their abundance. On the other hand, we find that proteins on other chromosomes show a surprising degree of differential expression. We propose that a large fraction of these differential expression events might serve as a compensatory purpose by binding aggregation prone proteins upregulated due to their location on the aneuploid chromosome: We observe that up to 40% of the typically hundreds and sometimes more than 1000 differentially abundant proteins physically interact either in a complex or in a binary manner with their partners on the aneuploid chromosome. Given the still incomplete understanding of the nature of the human interactome (and in particular the limits on the available protein complex information), this is a remarkably high number. We expect that with an increase of protein complex measurements, this number will substantially grow.

This novel compensatory mechanism complements the previously described dosage compensation addressing the differential expression of complex subunits directly on the aneuploid chromosomes^5,10^. Together they might largely prevent the otherwise detrimental overexpression of orphan complex subunits. Correlated abundance patterns to compensate for the differential expression of aggregation-prone proteins is only detectable for chromosome amplification cases (as it is expected as those are more likely to produce orphan subunits), while for deletions we observed stronger enrichments for specific, cancer-essential functions suggesting that here functional selection is a stronger driving force to shape the global co-abundance pattern.

Similar compensatory mechanisms have been previously identified to be induced by focal CNAs^8,9^. We demonstrate here that this observation holds for large genomic amplifications of entire chromosomes and likely serves the prevention of proteotoxic aggregation of orphan subunits as suggested by the stronger abundance correlations formed by aggregation-prone proteins. Considering that around 90% of solid tumors are aneuploid, our work addresses the question of how the vast gene expression changes induced by the amplification of large genomic regions can be tolerated by the majority of cancer cells.

One surprising observation is the presence of strong negative correlations between aneuploid proteins and co-complex partners on other chromosomes. We could not substantiate our initial intuition that those could be indicative of protein binding competition relationships. We used different approaches to predict overlap in protein binding interfaces to estimate competition events but did not observe an agreement with the negative abundance correlations. Future research will need to clarify the reason for the existence of the negative correlations.

Taken together, our findings describe the need for compensation mechanisms to deal with stoichiometric imbalances in protein complexes induced by aneuploidy, highlight their clinical relevance and shed light on the underlying mechanisms. Ultimately, given the high number of aneuploid tumors, studying and understanding compensatory mechanisms and the potential vulnerabilities they create in aneuploid tumors will have profound implications for both basic cell biology as well as cancer biology.

## Materials and methods

### Calculating whole-chromosome level aneuploidy scores

Arm-level aneuploidy scores for 10,522 TCGA samples, comprising 33 cancer types, were obtained from Taylor et al^4^. Whole-chromosome level aneuploidy scores were calculated as follows: If both p and q arms for chromosomes 1-12 and 16-20 are amplified, deleted or not changed, the entire chromosome was considered as amplified, deleted or diploid, respectively.

For acrocentric chromosomes, 13-15 and 21-22, q arm aneuploidy scores were considered as representative for whole chromosome-level aneuploidy scores (Supplementary file 1). TCGA samples that have conflicting events (amplification, deletion or no change) on different arms or missing data for one or both arms were removed from further analyses. In this study, colon (COAD) and rectum adenocarcinoma (READ) were considered as one cancer type as COREAD.

### Detecting cancer type-specific whole chromosome-level aneuploidies

Chi-square test was performed to test the occurrence of a whole chromosome-level aneuploidy within each cancer type against random expectation. Then multiple testing correction was applied on the p-values by using Holm’s method. Cancer type-specific whole chromosome-level aneuploidies were selected based on the criteria that adjusted p-value lower than or equal to 0.05 and chi-square standard residual equal to or higher than 2, resulting in 86 and 117 whole chromosome-level amplifications and deletions, respectively (Supplementary file 1).

For each of the 86 cancer type-specific amplifications, co-amplification frequency with other chromosomes were tested by using chi-square test. Then multiple testing correction was applied on the p-values by using Holm’s method. 305 significant combinations were identified as co-amplified events based on the criteria that the adjusted p-value was lower than 0.01 (Supplementary file 1).

### Data processing

RNA-seq FPKM values for 11,007 TCGA samples, comprising 32 cancer types, were downloaded from the NCI Genomic Data Commons (GDC)^26^. Then FPKM values were converted to TPM values and primary tumor samples (n = 9830) were selected. Ensembl gene IDs were mapped to gene symbols based on the mapping obtained from ensembl BioMart (Human genome version GRCh38.p13 - downloaded on May, 2019)^27^ and the mean value was taken when multiple Ensembl IDs mapped to one gene symbol. Mitochondrial and non-expressed (zero values in all samples) genes were removed.

Proteomics data used in this publication were generated by the Clinical Proteomic Tumor Analysis Consortium (NCI/NIH). Proteomics measurements for the available TCGA projects were downloaded from the CPTAC, covering 3 cancer types, spectral counts for COREAD^17,18^, and relative abundances for OV^15,16^, and BRCA^13,14^. For the replicated samples, the mean value was considered. Primary samples covered by the transcriptomic data and genes that are expressed at transcriptome level were selected, which gave us 88 samples and 5353 genes, 119 samples and 7062 genes, 105 samples, and 10467 genes for COREAD, OV and BRCA, respectively (Figure 1 – figure supplement 1A). Spectral counts for COREAD were normalized by quantile normalization followed by log2 transformation.

### Detecting transcriptomic and proteomic changes

To detect transcriptomic and proteomic changes, samples covered by aneuploidy, transcriptomic and proteomic data were selected, which resulted in 9266 samples for transcriptome analysis and 298 samples for proteome analysis.

For each of the 203 cancer type-specific whole chromosome-level aneuploidies covering 86 amplifications and 117 deletions, we first grouped TCGA samples as the ones with chromosome amplification/deletion and the ones diploid for the respective chromosome. After selecting the samples, genes having 0 TPM in all samples were filtered out. Differentially expressed genes between the samples with diploid and those with an altered chromosome were identified by using Wilcoxon test and then multiple testing correction was performed on the p-values by using the Benjamini and Hochberg method. Significantly differentially expressed genes were selected based on the criteria that the adjusted p-value is lower than 0.1. For the cases where we were left with less than 250 differentially expressed protein coding genes after adjusted p-value cutoff, the uncorrected p-value was used (p-value < 0.05) in order to have a sufficient number of genes to perform the enrichment tests (described below).

For the 13 cancer type-specific whole chromosome-level amplifications and 20 cancer type-specific whole chromosome-level deletions covering COREAD, OV, and BRCA cancer types, for which the corresponding proteome data is available, differentially abundant proteins between the samples with diploid- and amplified/deleted chromosome were detected by using Wilcoxon test. Proteins with a p-value lower than 0.1 were considered as significantly differentially abundant (again, using a relaxed statistical cutoff in order to perform the subsequent analyses of the protein set).

### Grouping proteins and protein pairs

Aggregation prone proteins (n=300) were obtained from Maata et al^22^. The known human protein complexes (n=2916) were downloaded from the CORUM database^19^ (CORUM 3.0, September 2018). The number of complexes a protein is involved in was calculated by considering the CORUM complexes, and then proteins were grouped as promiscuous if they are involved in more than 5 complexes, otherwise as non-promiscuous. To calculate co-occurrence frequencies, for each protein pair, we first counted the number of complexes in which the two proteins were found together and then divided it by the number of complexes in which at least one of them is found.

### Statistical analyses

Chi-square test was used to assess the relationship between differentially abundant proteins on other chromosomes and (i) complex members obtained from the CORUM database, and (ii) co-complex members of differentially abundant proteins on aneuploid chromosomes for the 13 cancer type-specific amplifications and 20 deletions. The same association tests were repeated by using transcriptome level changes - differentially expressed genes on other chromosomes - for 86 amplifications and 117 deletions, and then multiple testing correction was performed (only for transcriptome level analysis as here a much larger number of tests was performed) on p-values by using Holm’s method.

Cancer type-specific protein abundance correlations were calculated between all possible pairs across primary tumour samples for the cancer types BRCA, COREAD, and OV (for which we have proteomic measurements) by using the Spearman method. Comparing correlations between co-complex members and non-complex members was done by considering differentially abundant proteins on aneuploid chromosomes and their correlations with all CORUM complex subunits. The Wilcoxon test was used to compare the correlation distributions. To this end, correlations from 3 cancer types were pooled.

To compare the protein level correlations between different protein feature groups, correlations between differentially abundant proteins on aneuploid chromosomes (the ones that showed an increase in abundance for amplifications or a decrease for deletions) and their co-complex members were considered. To obtain a unique set, correlations from 3 cancer types were pooled. For the pairs for which we could compute a correlation in more than one cancer type, the maximum correlation value was considered, which left us with 2772 and 3818 correlations for amplifications and deletions, respectively (Supplementary file 4).

### Network randomization

To assess if there is an enrichment between differentially abundant proteins encoded on the aneuploid and those on other chromosomes, we employed a randomization test. We retrieved PPI data from HIPPIE (v2.2)^20^ and counted the number of physical PPIs between the two protein sets. In each randomization, we replaced the set of aneuploid differentially abundant proteins by a protein set of equal size. To avoid biases, we additionally enforced the same degree distribution to the original set by replacing each differentially abundant aneuploid protein by a protein of the same or similar degree (forming as many interactions as the replaced protein). We then recounted the number of PPIs between the random set and differentially abundant proteins encoded on other chromosomes to construct a background distribution from which we estimated the p-value by counting how often the original observed value was smaller or equal than a randomized value (Supplementary file 3).

### Functional annotation of protein complexes

To investigate functional relevance of co-abundance regulation, we first classified abundance correlations between differentially abundant proteins of aneuploid chromosomes and their co-complex members of other chromosomes into two groups: Top correlated ones including 20 strongest positive and negative correlations (40 in total) and non-correlated ones including correlations between -0.2 and 0.2. The latter was used as background in the association test. Then we obtained protein complexes and their functional annotations - associated gene ontology (GO) terms - from the CORUM human protein complex data. For each GO term, a chi-square test was performed in which the number of complexes related to the corresponding term in the top correlated group was tested against that of in the background group. An enrichment score was calculated by dividing the difference of the observed complex number and the expected one obtained from the chi-square test by the square root of the expected value. GO terms with p-value lower than 0.05 were considered as significantly associated. The analysis was done separately for each detected cancer type-specific aneuploidy case (13 amplifications and 20 deletions covering BRCA, OV, and COREAD cancer types for which we have proteomic measurements).

### DNA methylation analysis

Promoter level methylation measurements calculated from probe level methylation data in TCGA was used in this analysis^28^. For each gene, the most upstream promoter was considered for the analysis. Average methylation level of genes was calculated by taking the mean of methylation levels across aneuploid samples and diploid samples, separately. Wilcoxon tests were performed for the statistical comparison between up- and downregulated genes.

### TF - target randomization test

To test if differential expression of TFs on aneuploid chromosomes could explain the vast expression changes on other chromosomes, we performed a randomization test for 203 detected cancer type-specific aneuploidies. We first retrieved 1651393 gene-TF associations detected by ChIP-Seq experiments from the Harmonizome database^23,24^ and then counted the number of targets of differentially expressed TFs on aneuploid chromosomes among differentially expressed genes on other chromosomes. To assign a background distribution, we recounted the number of targets for random TF sets for 100 iterations. In each iteration, the random TF set size was equal to the number of differentially expressed TFs on the aneuploid chromosome of the corresponding cancer type-specific aneuploidy case. The p-value was calculated using the background distribution by conducting a two-tailed test.

### Ubiquitination analysis

Experimentally observed ubiquitination sites for human proteins were downloaded from PhosphoSitePlus^25^. The unique set of abundance correlations between differentially abundant proteins of aneuploid chromosomes and their co-complex members of other chromosomes was used for this analysis (See Methods section: Statistical analyses; Supplementary file 4). For each co-complex member protein, the total number of ubiquitination sites was calculated as the sum of all ubiquitination sites. For the proteins not found in the ubiquitination dataset, the number of ubiquitination sites was assigned to zero. Abundance correlations equal to or higher than 0.4 and lower than or equal to -0.4 were considered as top positive and top negative correlations, respectively. Wilcoxon test was used for statistical comparison.

### Calculation of the stoichiometry deviation score and survival analysis

To calculate the stoichiometry deviation score for each TCGA sample, the top 30 strongest tissue-specific correlation pairs (aneuploid protein and its partner) in amplification cases were taken. For each pair, a linear regression model was performed in which protein abundance of partner protein was dependent variable, and that of aneuploid protein was independent variable. Then the stoichiometry deviation score of a sample was calculated as the mean of absolute residuals in the regression models.

To perform survival analysis, we first grouped samples based on their stoichiometry deviation scores by using the survminer package (version 0.4.8) in R. Then, we performed survival analysis by using Kaplan Meier method in the survival package (version 3.1.8) in R.

GO annotations were retrieved from the UniProt^29^ database, and proteins with GO terms related to proteasome complex and ubiquitin-binding were selected. Then, the correlation between protein abundances and stoichiometry deviation scores was calculated across samples by using the Spearman method. Wilcoxon test was used for the comparison.

## Availability of data and materials

The code performing all analyses in this study is available at https://github.com/SengerG/Coregulation-of-complexes-in-Aneuploidtumors.git.

## Acknowledgements

We would like to thank Jason M. Sheltzer for his careful reading of our manuscript and very helpful comments.

## Supplementary Figures

**Figure 1 – figure supplement 1.**
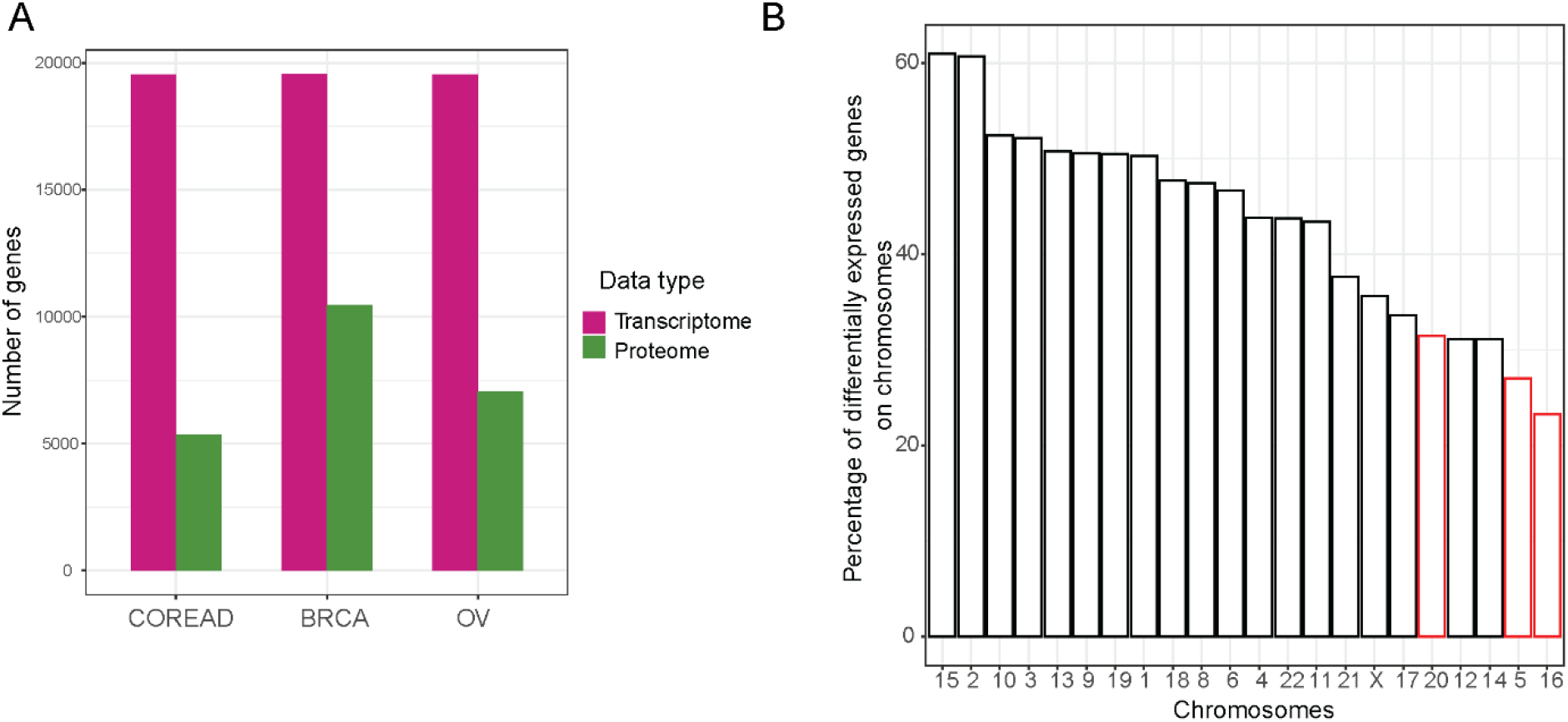
Proteome data coverage and differential expression changes on other chromosomes. **A)** The number of genes covered by transcriptomic and proteomic data for TCGA cancer patients comprising COREAD, BRCA, and OV cancer types. **B)** The effect of co-amplified chromosomes on the differential expression changes induced by THCA chromosome 7 amplification. Percentage was calculated as the ratio of differentially expressed genes from each chromosome to the total number of expressed genes on that chromosome. Red bars show the percentage for chromosome 5, 16 and 20 which are strongly co-amplified with chromosome 7 in THCA.

**Figure 2 – figure supplement 1.**
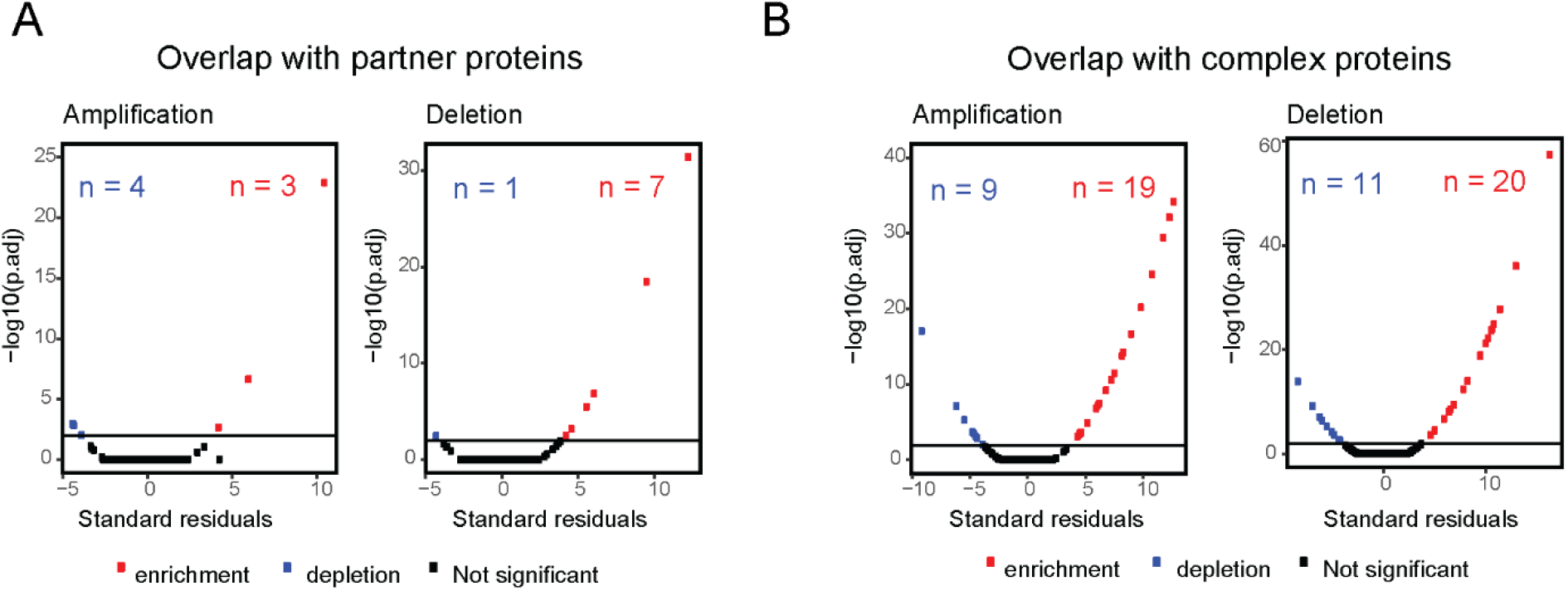
Transcriptome level changes on other chromosomes. Standard residuals and adjusted p-values for the overlap between **A)** co-complex members of differentially expressed genes on aneuploid chromosomes and differentially expressed genes on other chromosomes, for 203 detected cancer type-specific aneuploidies; 86 amplifications and 117 deletions, and **B)** CORUM complex subunits and differentially expressed genes on other chromosomes.

**Figure 5 – figure supplement 1.**
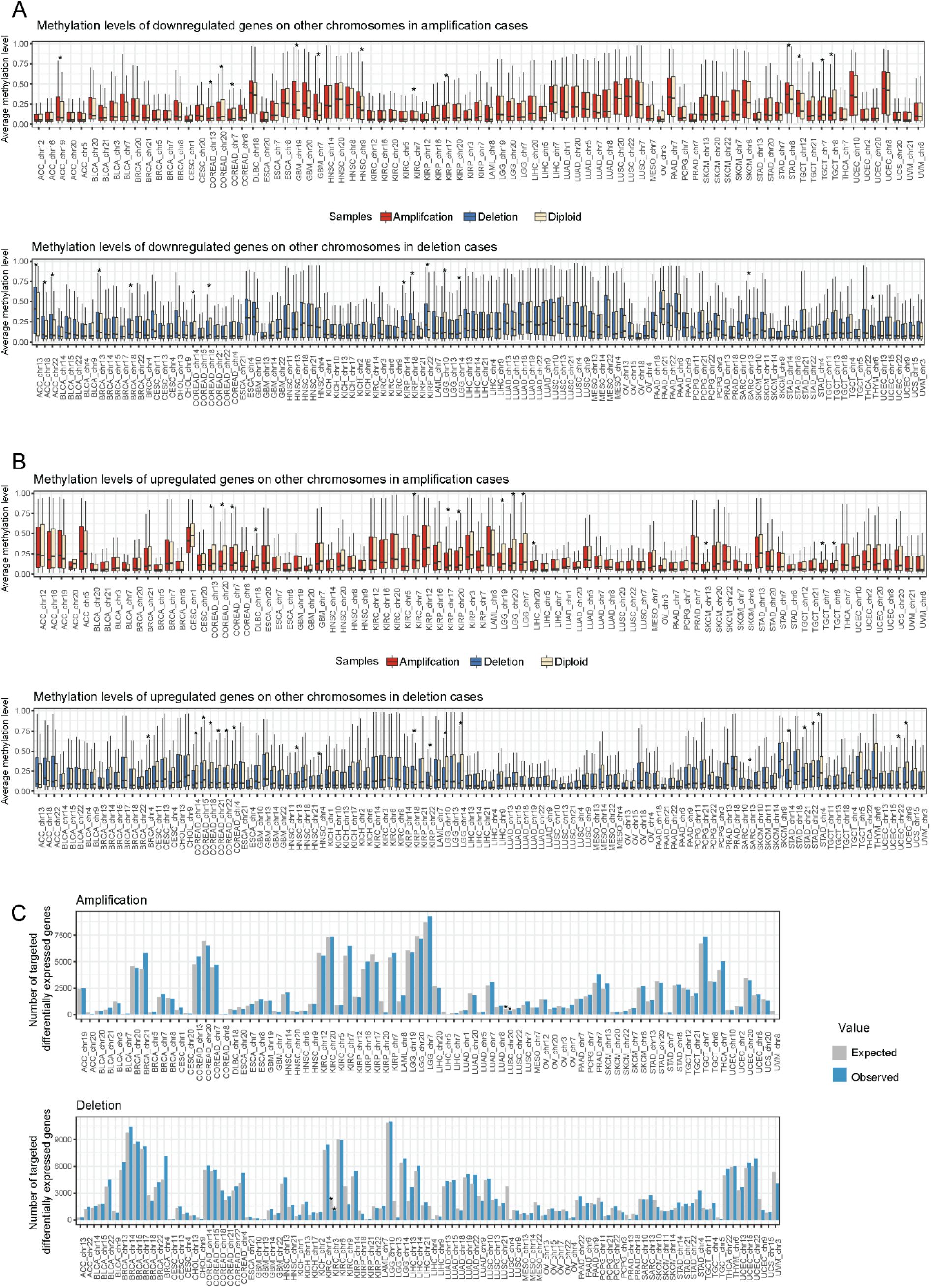
Expression changes on other chromosomes cannot be fully explained by transcriptional regulation. Average methylation level of **A)** downregulated and **B)** upregulated genes in aneuploid vs diploid samples. Wilcoxon test was used to test differences in methylation level of genes between aneuploid and diploid samples (*: P-value < 0.05). **C)** The number of targets of differentially abundant TFs on aneuploid chromosomes among differentially expressed genes on chromosomes. Expected value and p-value were calculated using a randomization test (*: P-value < 0.05).

**Figure 6 – figure supplement 1.**
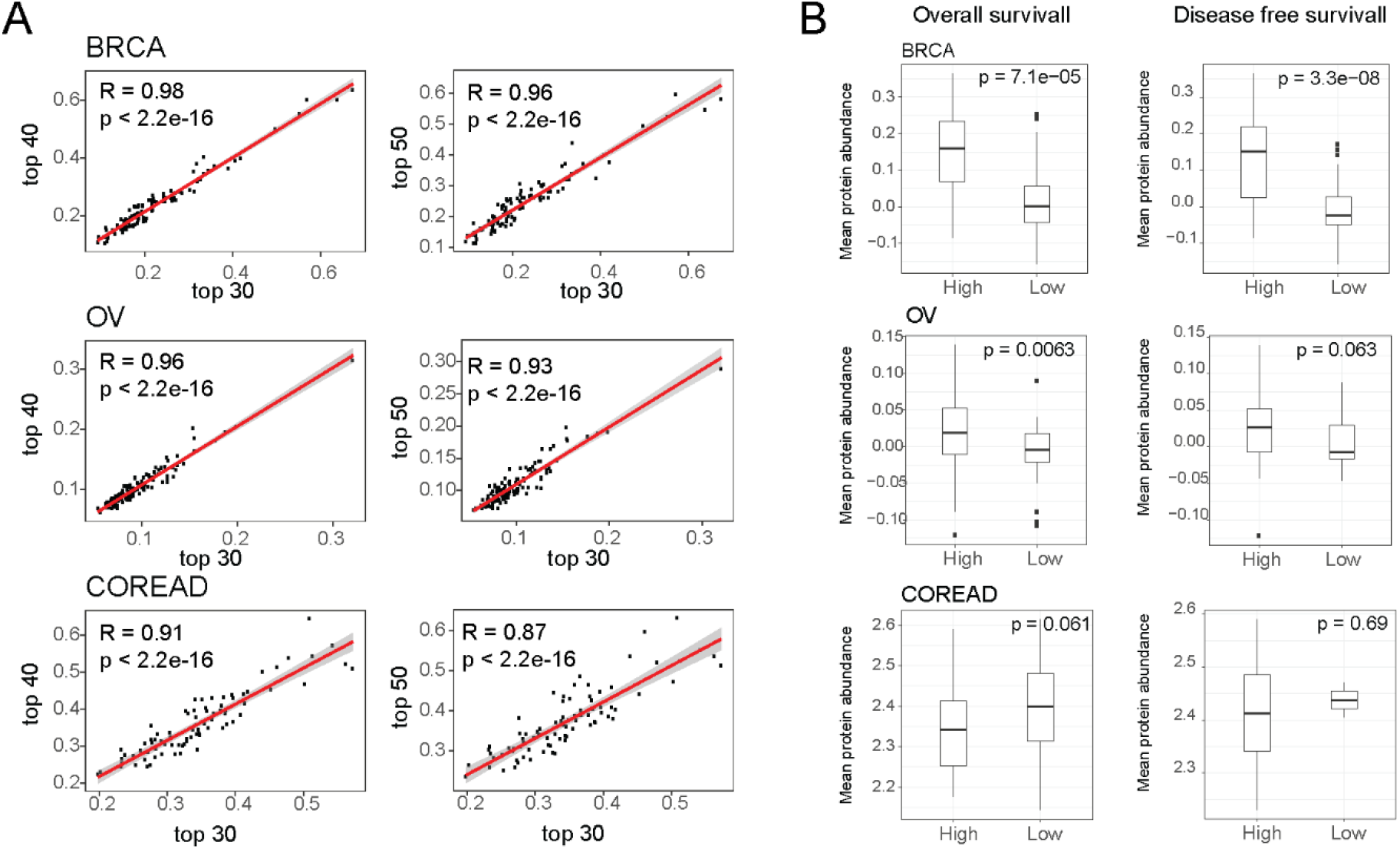
Association between the deviation from complex stoichiometry and survival probability. **A)** Correlations between the mean stoichiometric deviation scores across samples when different number of highly correlated protein pairs were considered. **B)** Mean abundances of proteasome complex and ubiquitin-binding proteins in high and low samples. Samples were grouped once based on overall survival, and once based on disease-free survival.

## Supplementary File Legends

**Supplementary File 1**. Whole chromosome-level aneuploidy scores, cancer type-specific whole chromosome-level amplifications and deletions, and co-amplified chromosomes, related to Figure 1 and Figure 1 – figure supplement 1B.

**Supplementary File 2**. The number of differentially abundant proteins on other chromosomes in partners of differentially abundant aneuploid chromosomes, in all expressed proteins, and in CORUM subunits. Standard residuals and p-values for the overlap between differentially abundant proteins on other chromosomes and partners of differentially abundant aneuploid proteins.

**Supplementary File 3**. Network randomization results; the number of PPIs between differentially abundant proteins on aneuploid chromosomes and those on other chromosomes and their corresponding p-values, related to Figure 2C.

**Supplementary File 4**. Protein level Spearman correlations between differentially abundant proteins on amplified/deleted chromosomes and their co-complex members on other chromosomes, and groups of proteins and protein pairs, related to Figure 3.

**Supplementary File 5**. Functional enrichment analysis of protein complexes; significantly enriched GO terms in different aneuploidy cases, their related p-values and enrichment scores, related to Figure 4.

